# Complex patterns of hitchhiking mutation load among stickleback populations

**DOI:** 10.1101/2025.07.24.666323

**Authors:** Jana Nickel, Jan Laine, Andrew D. Foote

**Author notes:** Corresponding authors: Centre for Ecological and Evolutionary Synthesis, Department of Biosciences, University of Oslo, 0316 Oslo, Norway.

## Abstract

Positive selection causes beneficial alleles to rapidly rise to high frequency in a population. This can cause linked genetic variation to **"**hitchhike**"**, and thereby also rise in frequency. This linked variation may include deleterious recessive alleles, previously neutrally harbored at low frequency in heterozygous genotypes. Modeling studies have shown that local effective population size and reduced recombination rate should contribute to the probability of deleterious mutations being swept to high frequency by being linked to beneficial alleles in selective sweeps or inversions. Marine sticklebacks have repeatedly adapted to thousands of freshwater habitats that became available after the last ice age, resulting in the formation of distinct marine and freshwater ecotypes. Selection acts on ancient standing genetic variation present in marine populations, causing freshwater-adaptive alleles to increase rapidly over tens of generations. These genomic regions play a key role in the repeated freshwater adaptation of morphological, physiological, and behavioral traits, and evolve under strong selection. Thus, threespine stickleback are an ideal system for investigating the impact of hitchhiking mutation load. We estimate the mutation load in regions of low recombination, including inversions, and the *Eda* haplotype. We find little evidence for the accumulation of deleterious alleles among genomic regions and populations. Inversions deviated from Hardy-Weinberg equilibrium in several populations, exhibiting an excess of homozygotes. These observations are consistent with purging facilitated by recombination in inversion karyotypes. We conclude that adaptation in threespine stickleback is not limited by hitchhiking deleterious mutations.

**Significance statement:** Hitchhiking is a ubiquitous force in population genetics, shaping genetic diversity across the genome. Threespine stickleback are a model system for studying parallel adaptation to ecological variation, with low recombination genomic regions known to drive ecological adaptation. A key question is whether a deleterious mutation load is associated with adaptation in this system. We find that hitchhiking mutation load patterns are complex and do not conform to simple model predictions, despite some widespread patterns shared among populations. Our findings suggest that purging in homozygous karyotypes may be critical in reducing deleterious mutation load.

## Introduction

Felsenstein (1) elegantly argued that the evolutionary advantage of recombination was to disassociate linkage between beneficial and deleterious alleles, and to link adaptive alleles on the same haplotype. The relationship between recombination and linkage of beneficial and deleterious alleles on a haplotype predicts that regions of low recombination rate in the genome may have a reduced capacity to disassociate linked beneficial and deleterious mutations. Low recombination regions include those evolving under strong linked selection (e.g., selective sweeps), or within structural variants such as inversions (2, 3).

The importance of structural variants within genomes in determining phenotypic variation and underpinning adaptations associated with divergent ecological or sexual selection within wild populations is of key interest to evolutionary biologists (4, 5). Inversions in a heterozygous state (‘heterokaryotypes’) suffer from recombination suppression and therefore a reduction in cross-overs between the two forms of the inversion. This can have an additional effect of linking many haploblocks into a single so-called ‘supergene’, in which many ecology- or phenotype-associated alleles across multiple loci can become linked on an inverted haplotype (6). Inversions have been linked to trait variation, including the striking phenotype differences of mating morphs of ruff (*Philomachus pugnax*) (7); tail length and coat color differences between prairie and forest ecotypes of deer mice (*Peromyscus maniculatus*) (8), and seed size and flowering time variation between prairie and dune ecotypes of sunflowers (*Helianthus* spp.) (9, 10). In the latter examples, ecological selection has repeatedly acted on the genetic variation within inversions resulting in parallel evolution of ecotypes and driving genetic divergence between them (8, 11). The role of inversions in behavioral, ecological, and phenotypic divergence is thought to maintain inversion polymorphisms (12).

Deleterious mutation load has been found to be associated with inversions in a range of taxa (*Heliconius numata* (13); *Acropora kenti* (14)). However, few empirical studies have focused on deleterious recessive mutation load associated with selective sweeps, despite both theoretical expectations (15) and empirical evidence (16) that mutation load can interfere with and slow the progress of adaptation. In this study, we look for empirical support for these model-based hypotheses (15) in natural populations of threespine stickleback with different demographic histories and consequentially effective population sizes. We compare mutation load in genomic regions with reduced recombination rate with putatively neutrally evolving regions.

Threespine stickleback provide an excellent system for understanding parallel adaptation to ecological variation (17). Marine stickleback have independently adapted to thousands of freshwater habitats that became available after the Last Glacial Maximum, resulting in the formation of distinct marine and freshwater ecotypes. Upon colonization of new freshwater habitats, selection acts on ancient standing genetic variation (SGV) present in marine populations, causing freshwater-adaptive alleles to increase rapidly over tens of generations (18–20). Quantitative trait loci (QTL) within some of these divergent genomic regions have been characterized and found to drive the behavioral, morphological, and physiological differences between ecotypes, such as the number of bony plates and body shape (21). These QTL include the Ectodysplasin (*Eda*) haplotype on chromosome IV, a highly pleiotropic locus linked to multiple traits that characterize differences between marine and freshwater sticklebacks, including bony plate architecture (19), lateral line patterning (22), and schooling behavior (23). The *Eda* gene exhibits a large selection coefficient and a large fitness effect (24). Consequently, freshwater ancestry rapidly increases at the *Eda* haplotype within a few generations in transplant experiments (24, 25), and after recent colonization events in natural lakes (26). Strong selection results in the 16 kb long genomic region of the *Eda* haplotype being shared by freshwater populations (19). This is contrasted with the high divergence up- and downstream of the flanking breakpoints, as alternative haplotypes are fixed in different lakes (27, 28), consistent with a selective sweep. Given the characteristics and strength of selection acting on the *Eda* locus during the colonization of freshwater, we predict that deleterious recessive alleles could hitchhike upon the *Eda* locus during the resultant sweep (**Figure 1**).

**Figure 1.**
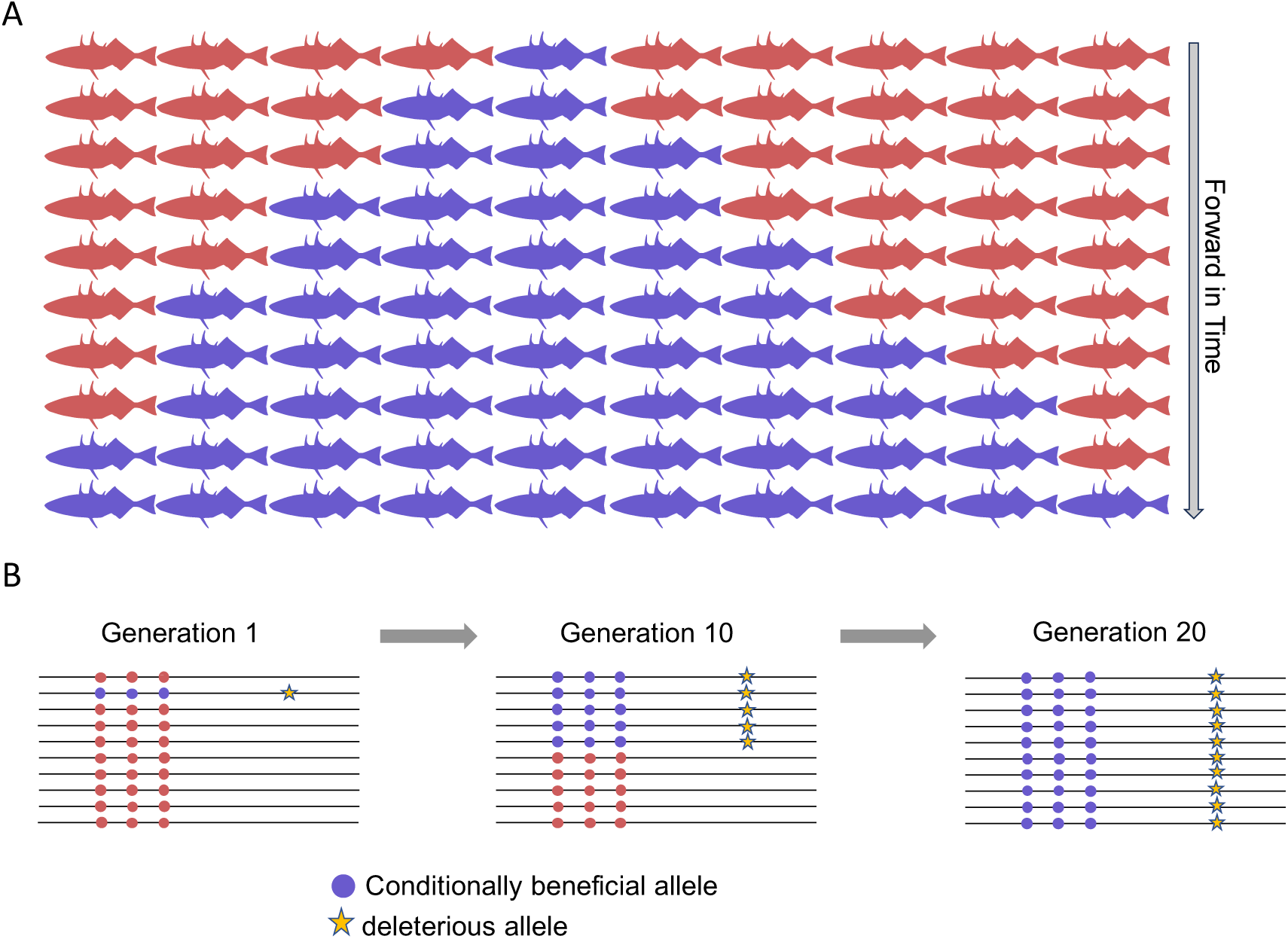
Schematic figure depicting **A.** changes in frequency of the freshwater haplotype (blue) and marine haplotype (red) every two generations for the first 20 generations after colonization of freshwater by a marine founder population. **B.** increase of conditionally beneficial freshwater alleles (blue) over generations and linked increase of deleterious alleles (yellow star). Stickleback silhouette: phylopic, Milton Tan.

The *Eda* locus is one of the 242 genomic loci, including several inversions, that are known to be associated with parallel marine-freshwater divergence (29). These marine-freshwater-adaptive loci typically display reduced recombination rates, thus maintaining strong linkage among loci on the same chromosome and creating **"**adaptive cassettes**"** that are under selection (30). Thus, we can compare the accumulation of deleterious mutations between inversions and within the selective sweep at the *Eda* haplotype to better understand the interaction between genomic architecture and mutation load.

## Results

We sequenced 241 stickleback from two assumed anadromous populations from marine fjords carrying predominantly marine-adaptive alleles at marine-freshwater divergent regions of the genome (see Jones et al. (29)), six lacustrine populations carrying predominantly freshwater-adaptive alleles, and one riverine population with mixed marine and freshwater ancestry (31), yielding an average genome coverage of 14.45 ± 6.23× (**Table S2**).

### Demographic history

We reconstructed the demographic history of the marine or freshwater populations, as effective population size (*N_e_*) determines the efficacy of selection. We inferred *N_e_* over time with SMC++. Assuming a generation time of 2 years (32), the maximum effective population size was reached 100,000 years before present (BP) (**Figure 3, Figure S4**). The two marine populations, Oslofjord and Altafjord, remained relatively stable over time with an effective population size of around 20,000. Freshwater populations experienced a decline in *N_e_* around 10,000 years BP, in particular the three oldest lakes, Langevatnet, Klubbvatnet, and Jossavannet, and exhibit lower effective population sizes. This demographic history would be consistent with the freshwater populations being isolated after the last ice age (27, 32). Due to the limited number of recent coalescent events, estimates of *N_e_* closer to the present should be interpreted with caution (33).

### Shared *Eda* haplotypes among lakes

We first focused on the selective sweep at the region containing the *Eda* gene and the flanking regions on chromosome IV to compare genomic differentiation and divergence between marine and freshwater populations. Overall, genome scans comparing freshwater populations and marine populations showed a similar pattern for all pairwise comparisons, with differentiation (*F*_ST_) and divergence (*d*_xy_) peaks at the *Eda* gene and the trailing ∼40 kb genomic region, followed by a smaller peak in the flanking region (**Figure 2A&B, Figure S1**). Although the two marine populations are approximately 1,350 km apart, they show very little genomic differentiation (*F*_ST_ = 0.01) and divergence (*d*_xy_ = 0.004) across the genome. Thus, the two marine populations can be used interchangeably.

**Figure 2.**
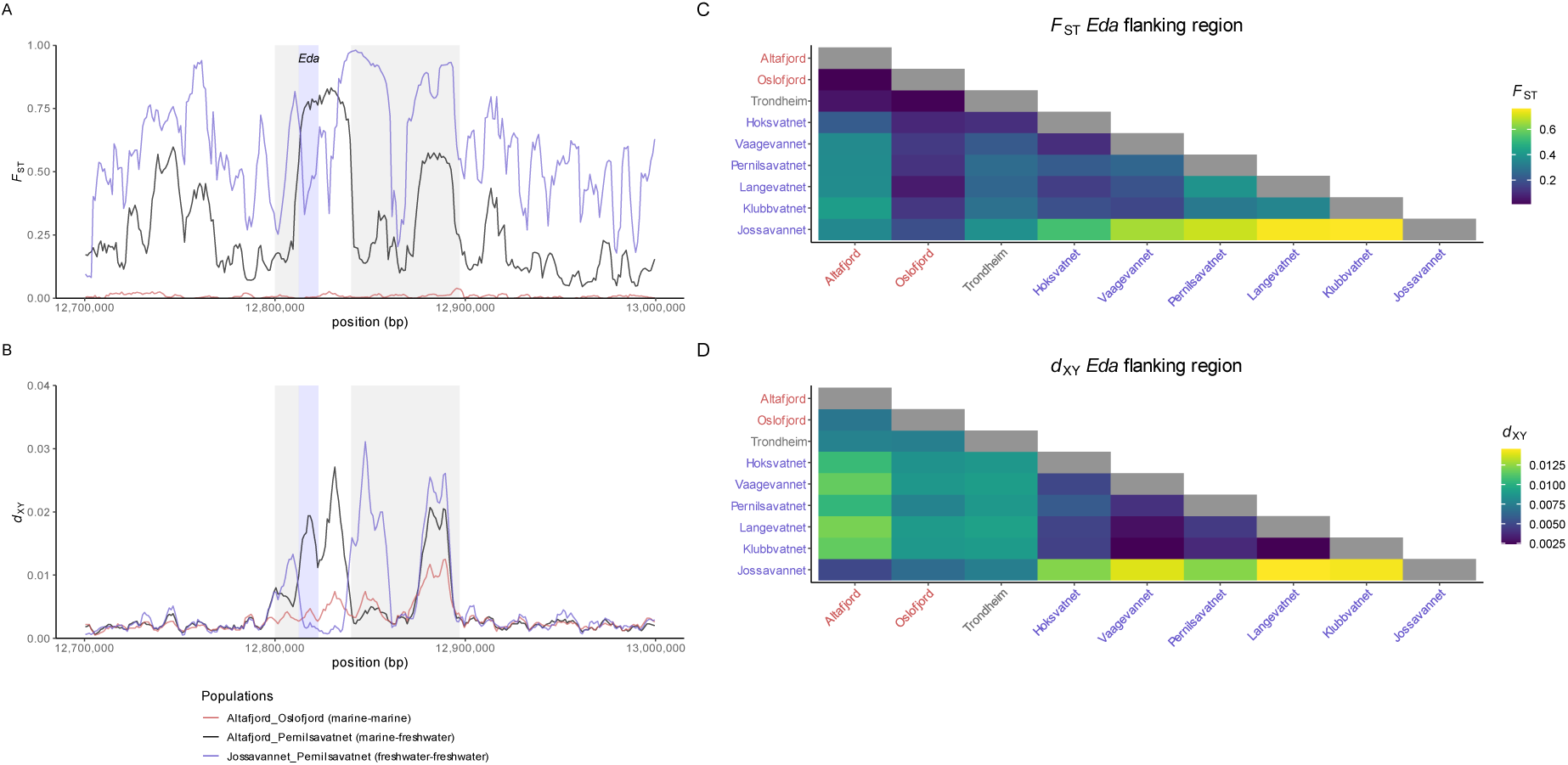
Window-based **A.** F_ST_ and **B.** d_xy_ statistics for exemplary population pairs Altafjord-Oslofjord (marine-marine), Altafjord-Pernilsavatnet (marine-freshwater), and Jossavannet-Pernilsavatnet (freshwater-freshwater) in 5 kb windows with 1 kb step size, showing the 300 kb genomic regions containing the *Eda* haplotype on chromosome IV. The location of the *Eda* gene is marked in blue. Based on the peaks, we defined the surrounding upstream and downstream flanking regions (grey). All population pairs are shown in **Figure S1**. Window-based **C.** *F*_ST_ and **D.** *d*_XY_ statistics for all population pairs averaged for the combined upstream and downstream flanking regions (47 genomic windows) as a heatmap. Marine populations are written in red and freshwater populations in blue. Separate *F*_ST_ and *d*_XY_ statistics for the upstream and downstream flanking regions are shown in **Figure S3**.

For most freshwater-freshwater population comparisons, we find reduced divergence at the *Eda* locus and the trailing genomic region with peaks in the flanking regions, i.e., differentiation is reduced directly at the *Eda* locus but is relatively high in adjacent regions of chromosome IV (**Figure S1**). This pattern more broadly supports the previously reported findings of reduced *F*_ST_ at the *Eda* locus and high *F*_ST_ at the flanking regions in comparisons between freshwater populations from Vancouver Island (28) and between the Jossavannet and Klubbvatnet populations (27). Comparisons between marine or freshwater populations reveal genetic differentiation and divergence peaks at the *Eda* gene, while comparisons between freshwater populations show reduced differentiation and divergence. This suggests that the core *Eda* haplotype is shared by these six isolated lake populations.

We visualized the average *F*_ST_ and *d*_xy_ as a heatmap in the upstream and downstream flanking regions of the *Eda* gene, comparing all populations (**Figure 2C&D, Figure S3**). In the upstream and downstream flanking regions, the differentiation and divergence are lowest between the marine populations, and are most elevated in all comparisons that include Jossavannet, which is known to be highly inbred, with over 60 % of the genome being found in long (>300 kb) runs of homozygosity (27). These patterns are consistent with linked haplotypes adjacent to *Eda* approaching fixation in at least some freshwater populations, i.e., a hard selective sweep. All other comparisons between freshwater populations show slightly increased differentiation and divergence in the flanking regions. The nucleotide diversity is markedly reduced in most freshwater populations, while populations Altafjord, Oslofjord, Trondheim, and Hoksvatnet show a peak at the *Eda* gene and the downstream flanking region (**Figure S2**). Samples collected approximately 1 km upstream of the Nidelva River mouth for the Trondheim population had a mixture of marine or freshwater ancestry and traits and are therefore not categorized as either a marine or freshwater population in subsequent analyses.

### Mutation load in low recombination genomic regions and differences between populations

Our goal was to identify whether there is evidence for deleterious mutations segregating in low recombination regions, resulting in an increased mutation load. To test this, we compared the proportion of different categories of putative impact of variants in the selective sweep at *Eda* and the inversions on chromosomes I, XI, and XXI to the proportions of the genome-wide SNP data set (**Figure 4A**). The three inversions are associated with parallel marine-freshwater divergence (29), with chromosome XXI specifically harboring a high number of QTL (21).

**Figure 3:**
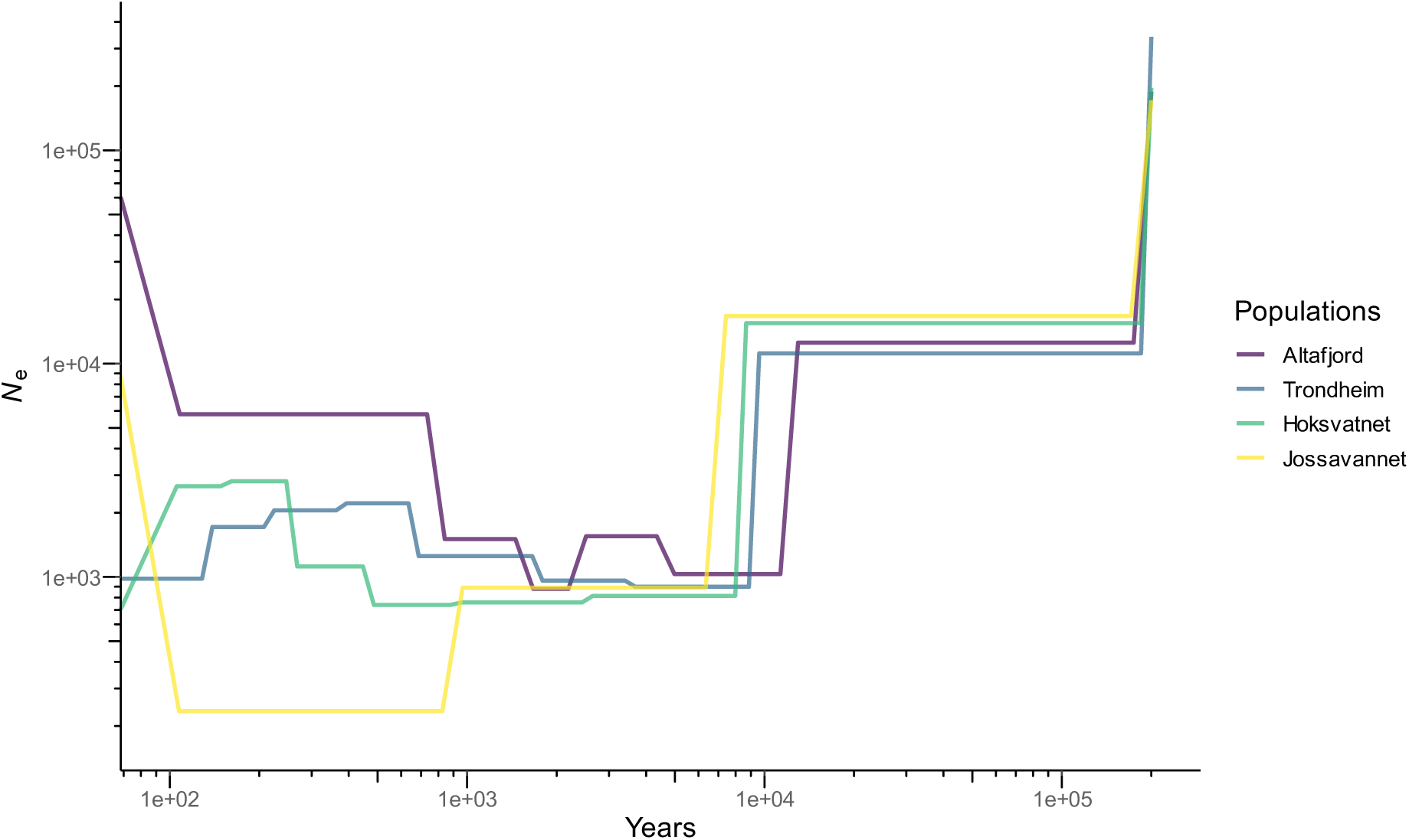
Demographic history for one marine (Altafjord), one river (Trondheim), and two freshwater populations (Hoksvatnet and Jossavannet) inferred with SMC++ using a generation time of 2 years and a mutation rate of 3.7 × 10^-8^ per site per generation. All populations are shown in **Figure S4**.

**Figure 4:**
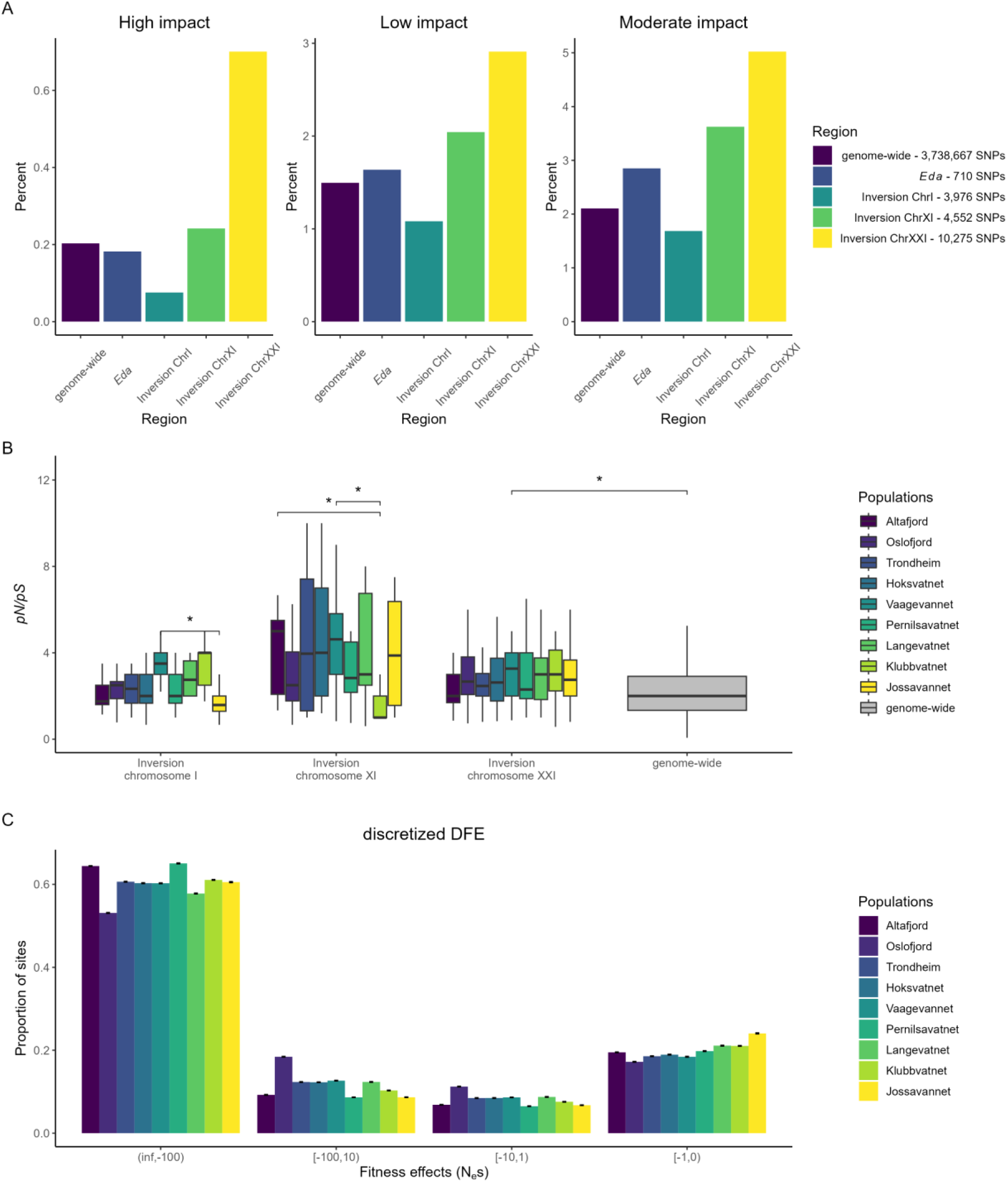
**A.** SnpEff estimated impact of SNPs for the whole genome (excluding the marine-freshwater divergent regions), the *Eda* region, and the three inversions. The legend shows the number of SNP annotations for each region. **B.** Ratio of nonsynonymous to synonymous sites (*pN*/*pS*) for the inversions on chromosome I, XI, and XXI for each population and genome-wide (excluding inversions) computed in 100 kb windows. The two-sided *t*-test showed that most comparisons between the populations for each inversion or between a population within an inversion and the genome-wide *pN*/*pS* were nonsignificant, and four comparisons were significant (*p* < 0.05). **C.** Discretized DFE estimate showing bins of population-scaled selection coefficient (γ = *N_e_*s) for each population. Error bars indicate 95% confidence intervals with 100 bootstraps.

For the whole genome, 0.2% of the SNPs were categorized as highly deleterious, which likely disrupts the gene function, 1.5% were categorized as low, and 2.1% as moderate impact. The 300 kb region surrounding the *Eda* haplotype is relatively small and therefore does not contain enough SNPs for each category for a meaningful statistical comparison. The inversion on chromosome I has a lower proportion for each impact category compared to the whole genome, consistent with purging of deleterious mutations. In contrast, the other two inversions show a higher proportion of deleterious mutations relative to the whole genome, especially the 2 Mb inversion on chromosome XXI. Thus, the relationship between inversions and mutation load is complex in threespine stickleback.

To characterize the mutation load of each inversion, we calculated the ratio of segregating nonsynonymous to synonymous mutations (*pN*/*pS*) in each region of interest of each population (**Figure 4B**) and for all marine or freshwater populations combined (**Figure S8**), and compared it to the genomic background. A higher ratio of nonsynonymous to synonymous mutations indicates potentially increased accumulation of deleterious alleles in a genomic region. We found no significant evidence of increased mutation load in any of the three inversions in marine or freshwater populations combined (**Figure S8**).

The two-sided *t*-test showed that the mutation load, estimated as *pN*/*pS*, was significantly different (*p* < 0.05) for the inversion on chromosome I between populations Jossavannet and Vaagevannet, and for the inversion on chromosome XI between populations Altafjord and Klubbvatnet, as well as between Klubbvatnet and Pernilsavatnet. The only value that was significantly different from the genome-wide *pN*/*pS* was the inversion on chromosome XXI for the population Vaagevannet (**Figure 4B**). The other comparisons between populations within each inversion and between a population within an inversion and the genome-wide *pN*/*pS* were nonsignificant.

For the distribution of fitness effects (DFE) in each population, we expect purifying selection to act more efficiently against highly deleterious mutations in large populations and be relaxed in smaller populations. In all populations, most SNPs are estimated to be deleterious, with most of them in the highest bin of population-scaled selection coefficient, indicating strongly deleterious mutations (**Figure 4C**). The proportion of sites with neutrally, weakly, moderately, and strongly deleterious fitness effects is similar across all marine or freshwater populations.

Regions with reduced recombination are expected to show increased mutation load due to less efficient purifying selection (1, 34). We compared the SFS for each population of different impact categories to identify potential purifying selection (**Figure S9**). The frequency of high, moderate, and low impact mutations for the whole genome, excluding the marine-freshwater divergent regions identified by Jones et al. (29), was not shifted compared to the neutral/modifier category in any population. The overall shape of the SFS suggests a history of population decline and bottlenecks.

### Inversion karyotype frequencies within and among populations

The standard karyotype, i.e., the most common form in the marine populations and therefore the expected ancestral state, is likely the cluster we assigned as A/A (**Figure 5**). The inversion karyotype is known to segregate between marine and freshwater populations (29) and is likely the cluster we assigned as B/B, which only appears in freshwater populations. In the marine populations, most individuals are homozygous for the assumed standard karyotype for all inversions. The highest inversion frequencies in freshwater populations are found for the inversion on chromosome I (57-100%), with fixation in the two oldest lakes, Klubbvatnet and Jossavannet. The majority of freshwater populations exhibit varying levels of polymorphism for all three inversions, which could point to differences in the evolutionary history of inversions.

**Figure 5:**
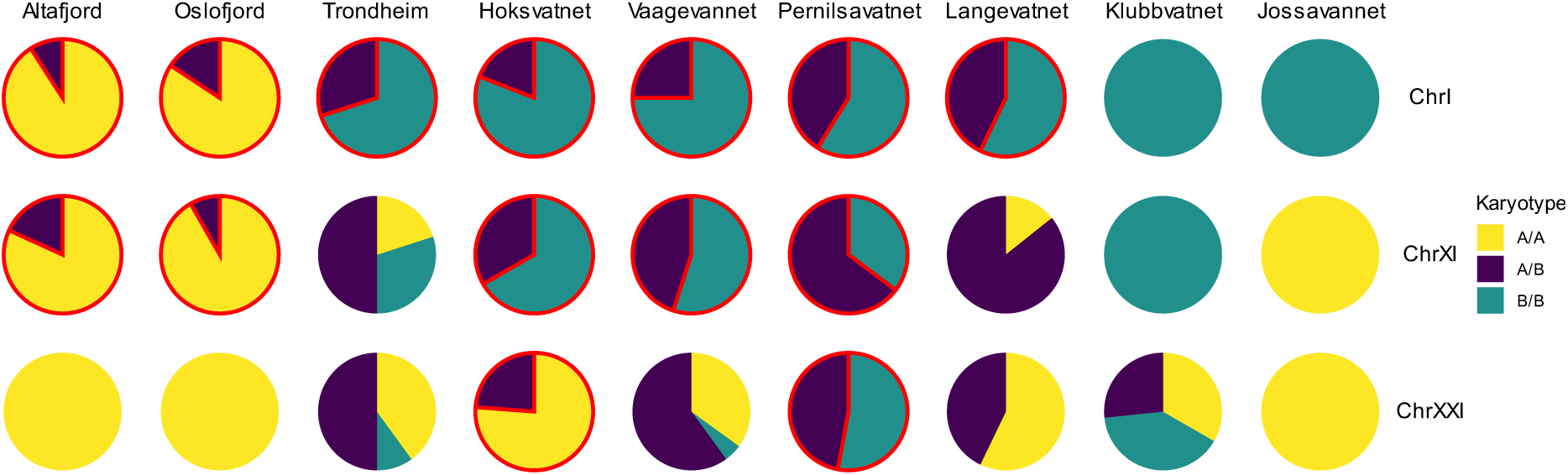
Distribution of inversion karyotype frequencies for all populations. Homozygous karyotypes are labeled as A/A and B/B, and the heterozygous karyotype is labeled as A/B. Populations that deviate from HWE are circled in red.

An alternative mechanism to reduce the impact of deleterious recessive alleles to purging is by associative- or pseudo-overdominance, in which recessive alleles are neutrally harbored in heterozygous genotypes. Populations varied in the degree of polymorphism and which karyotypes they carried at each inversion. The majority of marine or freshwater populations deviate significantly from Hardy-Weinberg equilibrium (HWE) at the inversions on chromosomes I and XI, while only two freshwater populations deviate from HWE for chromosome XXI. The excess of homozygotes in the marine and freshwater populations indicates strong spatially divergent selection.

## Discussion

Hitchhiking involves changes in allele frequencies linked to selective sweeps or to purged deleterious mutations (background selection), and is a ubiquitous force in shaping genetic variation in populations (35). The *Eda* locus is under strong selection in freshwater threespine sticklebacks, with a large selection coefficient (24) and effects on multiple traits (22). Due to the reduced recombination rate in selective sweeps, such as found at the *Eda* locus, we hypothesized that this region could have an increased hitchhiking mutation load. The freshwater threespine stickleback populations included in this study had distinct demographic histories due to being formed at different times following the Last Glacial Maximum. Colonization by marine stickleback was estimated to have occurred between 1,000 to 12,900 years ago for the different lakes sampled here. Demographic reconstructions show a decline in *N*_e_ around 10,000 years BP found in Atlantic stickleback (27, 32), and a larger effective population size in marine populations than in freshwater populations (**Figure 3**). Theory predicts that small populations should accumulate more deleterious mutations due to the stronger effect of genetic drift and relaxed purifying selection (36), a phenomenon that has been observed in natural populations (37–39). We found that the core *Eda* haplotype is shared by all six freshwater populations, regardless of lake age, as expected from strong selection on ancient genomic variation present in the marine populations that colonized freshwater habitats (40).

Despite the evidence for selection on the *Eda* haplotype in our freshwater populations, we were unable to confirm elevated mutation load due to the hitchhiking of deleterious alleles. This was due to a lack of power caused by the relatively small size of this genomic region and the limited number of SNPs carrying potential deleterious mutations. Quantifying *pN*/*pS* is only possible in coding regions of the genome and is more challenging in small genomic regions that contain only a limited number of synonymous or nonsynonymous mutations. Venu et al. (30) found that low recombination regions (or coldspots) are not limited to the flanking regions of the *Eda* haplotype, but also encompass a much larger region of linked genomic loci inherited together as a **"**linked adaptive cassette**"**. These regions shape marine-freshwater divergence and could be a target for investigating increased hitchhiking mutation load in low recombination genomic regions.

Inversions suppress recombination in their heterozygous form, thus resulting in increased linkage between alleles within inversions (4, 5). Inversions may also be affected by associative overdominance, where an inversion captures a karyotype carrying weakly deleterious, partially recessive, or recessive mutations at several loci that are masked in inversion homozygotes (5). This has been predicted to reduce purging of deleterious mutations from inversions, and to potentially promote associative overdominance, thereby limiting ecological adaptation (41).

Focusing specifically on three large inversions with some known association between marine and freshwater stickleback populations (29), we found inconsistent patterns of inferred deleterious mutation load in inversions. The largest inversion on chromosome XXI carries the highest percentage of high, low, and moderate impact SNPs compared to the genome-wide background and the other genomic regions (**Figure 4A**). The ratio of nonsynonymous to synonymous sites revealed that only the inversion on chromosome XXI in population Vaagevannet exhibited a significantly increased mutation load compared to the genome-wide background (**Figure 4B**). Chromosome XXI is known to harbor more QTL related to feeding, body shape, and defense than expected (21). However, lineage sorting of marine and freshwater populations at the inversion on chromosome XXI is less clear relative to the inversions on chromosomes I and XI (29). Thus, this inversion may have a limited role in marine-freshwater divergence.

Inversion frequency is shaped by balancing and divergent selection (42), and the tightly linked alleles are known to facilitate ecotype differentiation in fish (43–45), including stickleback (29). Inversion frequencies vary across the populations for chromosomes I, XI, and XXI, which could indicate differences in ecological association, evolutionary history, and inversion age (**Figure 5**) (5). All three inversions are highly polymorphic, but marine populations were monomorphic for karyotype A/A on the inversion on chromosome XXI, and the two oldest freshwater populations were mostly monomorphic. In freshwater populations, the inversion karyotype (B/B) is carried by the majority of individuals. In addition, the populations deviated from HWE with an excess of homozygotes, indicating positive selection on this karyotype in freshwater stickleback populations (46).

Theory predicts that reduced recombination rate in a genomic region increases the likelihood of deleterious hitchhiking alleles becoming fixed (15). However, this phenomenon appears to be more complex in natural populations and is influenced by several factors. A number of studies have identified an increased mutation load in inversions or supergenes, which also exhibit suppressed recombination. In the butterfly *Heliconius numata*, for example, three inversions involved in wing-pattern polymorphism were shown to have accumulated substantial deleterious mutations, but are maintained as polymorphic due to the reduced viability of homozygotes (13). Elevated mutation load has also been identified for multiple inversions in the coral *Acropora kenti* (14). In contrast, large inversions across multiple species of *Helianthus* exhibited a lower deleterious mutation load than expected. However, populations that were polymorphic for inversions exhibited a higher mutation load than monomorphic populations (47). Inversions play a role in the genetic differentiation and local adaptation of Atlantic herring, but do not carry increased genetic load. The authors suggest that this is due to the large population size, which allows for effective purifying selection (48). This is also the case for deer mice, where inversion homozygotes are common, and large effective population sizes could facilitate purifying selection (49).

Our study shares many of these key findings with the studies on deer mice (49) and sunflowers (47), whereby assortative mating or selection can result in deviation from HWE, with an excess of homozygote karyotypes. Deviation from HWE is common at inversions in the two marine populations and the younger freshwater populations. Our findings of mutation load in inversions showing limited differences to genome-wide patterns, suggest that recombination between homozygous inversion karyotypes is sufficient to purge excessive deleterious mutation load. However, the finding of fixation of the karyotype A/A (common in marine populations) at chromosomes XI and XXI in the most inbred freshwater population could support the hypothesis of Roesti et al. (41), that inversions can limit optimal ecological adaptation.

In a conservation context, mutation load in a population and its potential negative effect on fitness is often used to estimate the threat to small populations (50). In the Japanese stickleback, it has been shown that isolated freshwater populations accumulate more deleterious mutations than marine populations (51). Our study is among emerging studies that consider the potential impact of increased mutation load in low recombination genomic regions caused by hitchhiking deleterious alleles. Understanding the real-life fitness effects of such mutation load requires future experimental validation.

## Materials and Methods

### Sample collection

Adult threespine sticklebacks were collected using unbaited minnow traps or shore-based seine netting from two marine and seven freshwater populations in Norway. The pectoral or caudal fin was clipped and preserved in 95% ethanol, and the fish were released back, or sticklebacks were euthanized and frozen. The lakes ranged in age from 1,000 to 12,900 years old, and further information on all populations is provided in **Table S1**.

In addition, threespine sticklebacks were collected from Jossavannet and Altafjord in 2019 and 2021 as part of previous studies, as described in Kirch et al. (27) and Laine et al. (54).

### DNA isolation, library preparation, and sequencing

DNA was extracted from the fin tissue using the Qiagen DNeasy Blood and Tissue Kit following the manufacturer’s protocol. Libraries were prepared with the NEBNext Ultra II FS DNA Library Prep Kit for Illumina and sequenced using 150 bp paired-end reads on an Illumina NovaSeq 6000 at the Norwegian Sequencing Centre. Short-read data are deposited in the NCBI Sequence Read Archive under BioProject accession number PRJNA693136.

### Read alignment and variant calling

The quality of the reads was assessed with FastQC v0.12.1 (53). We trimmed adapters, Ns, and low-quality bases using AdapterRemoval v2.3.1 (54). Where necessary, poly-G tails were trimmed with fastp v0.23.4 (55). The trimmed reads were mapped to the stickleback reference genome gasAcu1-4 (available at https://datadryad.org/stash/dataset/doi:10.5061/dryad.547d7wm6t, (26)) using the BWA-MEM algorithm in BWA v0.7.17 (56). This reference genome originates from a freshwater population and includes improvements for the mitochondrial genome and the genomic region of chromosome VII containing the *Pitx1* gene. The BAM files were sorted, duplicates were marked, and files originating from the same samples were merged using samtools v1.16.1 (57).

Genotype calling was performed using the BCFtools v.1.20 (58) mpileup and call commands with the multiallelic calling model. To calculate the total number of genotyped sites (variant and invariant), we removed indels and multiallelic sites with BCFtools v.1.20 to produce an invariant data set including 444,582,088 sites.

Subsequently, we removed sites for which genotypes were missing for more than 30% of the individuals, a Phred quality score of <30, a minimum genotype depth of 10×, and a maximum genotype depth of 100× using VCFtools v0.1.16 (59). The final SNP data set included 1,470,232 sites across 241 samples.

### Genome scan

We estimated nucleotide diversity (π), genetic differentiation (*F*_ST_), and divergence (*d*_xy_) from the invariant data set using the Python script popgenWindows.py (github.com/simonhmartin/genomics_general release 0.4, last accessed May 2024) with a sliding 5 kb window and a step size of 1 kb, comparing all possible freshwater-marine, freshwater-freshwater, and marine-marine population pairs.

### Mutation load

To identify increased accumulation of deleterious mutations in genomic regions of interest, we used the stickleback reference genome gasAcu1-4 annotation to build a custom database for SnpEff v5.2c (60). We then annotated the SNP file with SnpEff to predict the putative impact of variants in the entire genome and the regions of interest. The SNPs were classified into the categories high (e.g., stop-gain or frameshift variant), low (e.g., missense variant or inframe deletion), moderate (e.g., synonymous variant), and modifier (non-coding variants or variants affecting non-coding genes).

To test for increased accumulation of deleterious mutations in inversions compared to the genome-wide background, we calculated the ratio of synonymous to nonsynonymous variants (*pN*/*pS*) using the SnpEff annotation. We filtered the SNP file for private SNPs for the respective population using BCFtools v.1.20 and calculated *pN*/*pS* in 100 kb nonoverlapping genomic windows for each population for the genome-wide background (excluding the inversions) and for the three inversions. We performed a two-sided *t*-test in R v4.3.2 to test for significant differences in *pN*/*pS* between populations, as well as between the inversions and the genome-wide background.

To analyze whether the karyotype frequencies of inversions in each population were deviated from HWE, which could indicate a deleterious effect of homozygotes, we identified the heterozygous and the two homozygous karyotypes in all individuals based on a principal component analysis (PCA) with *K* = 3. The heterozygosity within each inversion for each individual was calculated with BCFtools v.1.20. Based on this, we assigned the cluster in the center of the PCA and with the highest heterozygosity as the heterozygous karyotype (A/B), and the other two clusters as the homozygous karyotype (A/A and B/B) (**Figure S5, Figure S6, Figure S7**). We then tested for HWE with the HWE.chisq function from the genetics package in R.

### Demographic analysis

To estimate the demographic history of each population, we used SMC++ v1.15.2, which uses a sequential Markov coalescent model to infer population size history from unphased genomic data (33). The VCF files were converted to the format used by smc++ with the vcf2smc command. The model for the population size history was fitted for each chromosome using the smc++ estimate command with a mutation rate of 3.7 × 10^-8^ (32) and plotted with a generation time of 2 years.

### SFS inference

Using the SNP file annotated by SnpEff, we calculated allele frequencies at each site for the derived allele in each population using the Python script freq.py and the genome-wide 1D folded site frequency spectrum using sfs.py (github.com/simonhmartin/genomics_general release 0.4, last accessed May 2024). We then filtered the annotated SNP file for the four impact categories: high, low, moderate, and modifier, and repeated the previous steps to produce genome-wide site frequency spectra (SFS) for each category. For our genomic regions of interest, we filtered the annotated SNP file for the marine-freshwater divergent regions (29) and produced SFS for each impact category to compare these with the genome-wide SFS.

### Distribution of fitness effects

To define ancestral and derived alleles, we used whole-genome data from one individual each of outgroup species *Gasterosteus nipponicus* (NCBI accession: DRR072169)*, Gasterosteus wheatlandi* (NCBI accession: DRR013347), and *Pungitius pungitius* (NCBI accession: SRR11611426). Mapping and genotype calling were performed as described previously, and the VCF files were merged and filtered using the same parameters. To annotate the ancestral alleles in the VCF file, we used fastDFE v1.1.8 with the maximum likelihood model, specifying the outgroups and requiring at least 15 ingroups at a site (61).

A key part of understanding natural selection is the distribution of fitness effects (DFE), which describes the proportion of new mutations that are expected to be deleterious, neutral, and beneficial (62). This can be achieved by contrasting the SFS of putatively neutral (synonymous) and selected (nonsynonymous) sites (61). The DFE were estimated for the whole genome for each stickleback population using the GammaExpParametrization model with 10 iterations and 100 bootstrap replicates with fastDFE v1.1.8. The DFE results were summarized into four bins of population-scaled selection coefficients: neutral (0 > *N*_e_s > −1), weak (−1 > *N*_e_s > −10), moderate (−10 > *N*_e_s > −100), and strong (−100 > *N*_e_s).

## Acknowledgments

This work was funded by an ERC Consolidator Grant – EXPLOAD (101045346) awarded to ADF.

## Notes

### Competing Interest Statement

The authors have declared no competing interest.

