## Appendix for "Complex patterns of hitchhiking mutation load among stickleback populations"

**This PDF file includes:**

Figures S1 to S9

Tables S1

**Other supporting materials for this manuscript include the following:**

Table S2

### Supplementary Figures

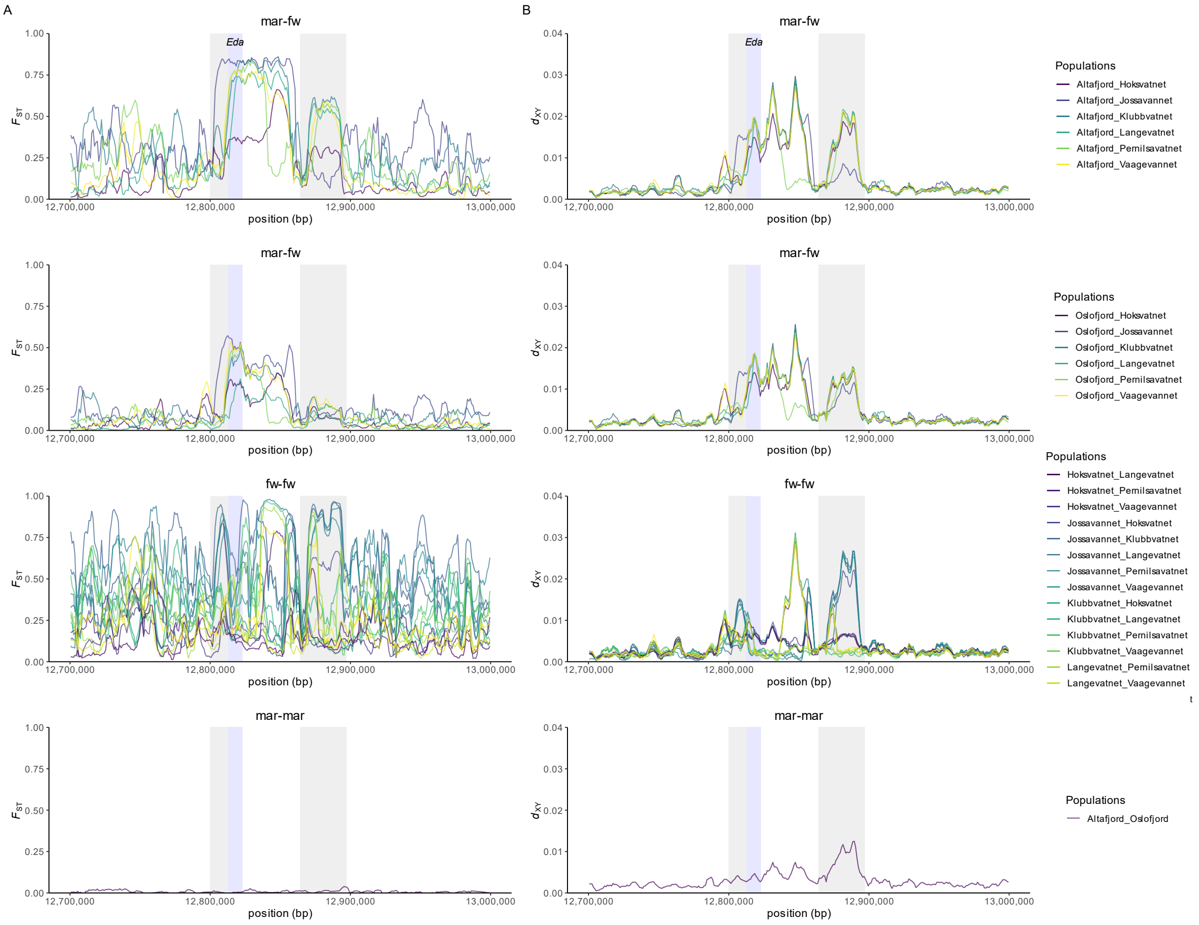


**Figure S1.** Window-based statistics for Altafjord-freshwater, Oslofjord-freshwater, freshwater-freshwater, and Altafjord-Oslofjord population pairs in 5 kb windows with 1 kb step size, showing the 300 kb genomic regions containing the *Eda* haplotype on chromosome IV. The location of the *Eda* gene is marked in blue. Based on the peaks, we defined the surrounding upstream and downstream flanking regions (grey). **A.** $F_{ST}$. **B.** $d_{xy}$.

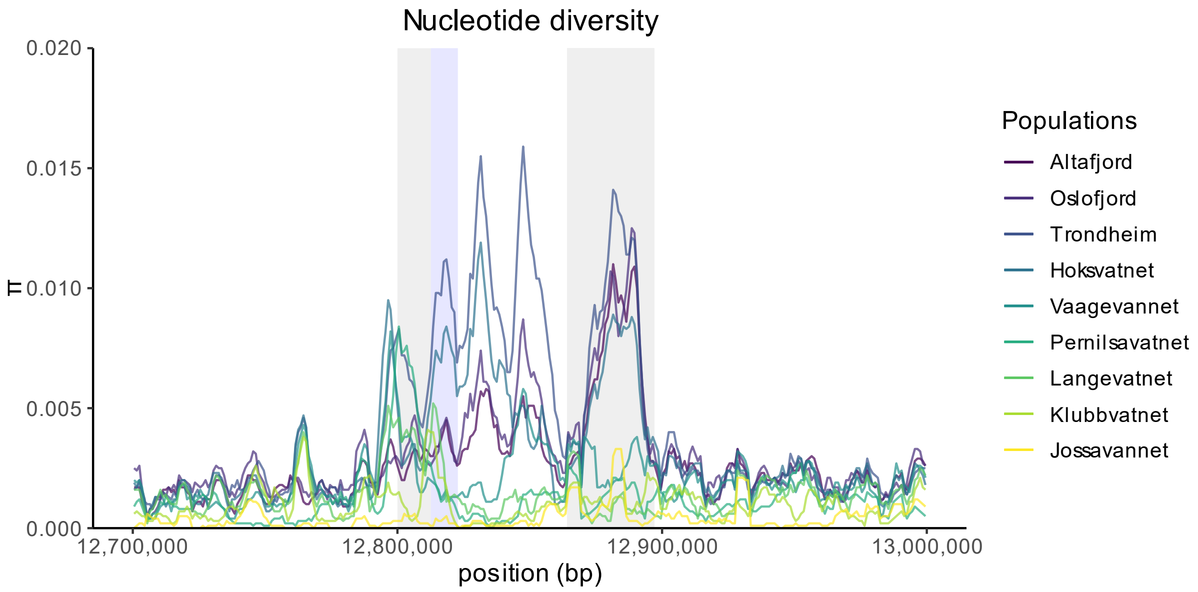


**Figure S2**: Window-based nucleotide diversity π for each population in 5 kb windows with 1 kb step size, showing the 300 kb genomic regions containing the *Eda* haplotype on chromosome IV. The location of the *Eda* gene is marked in blue.

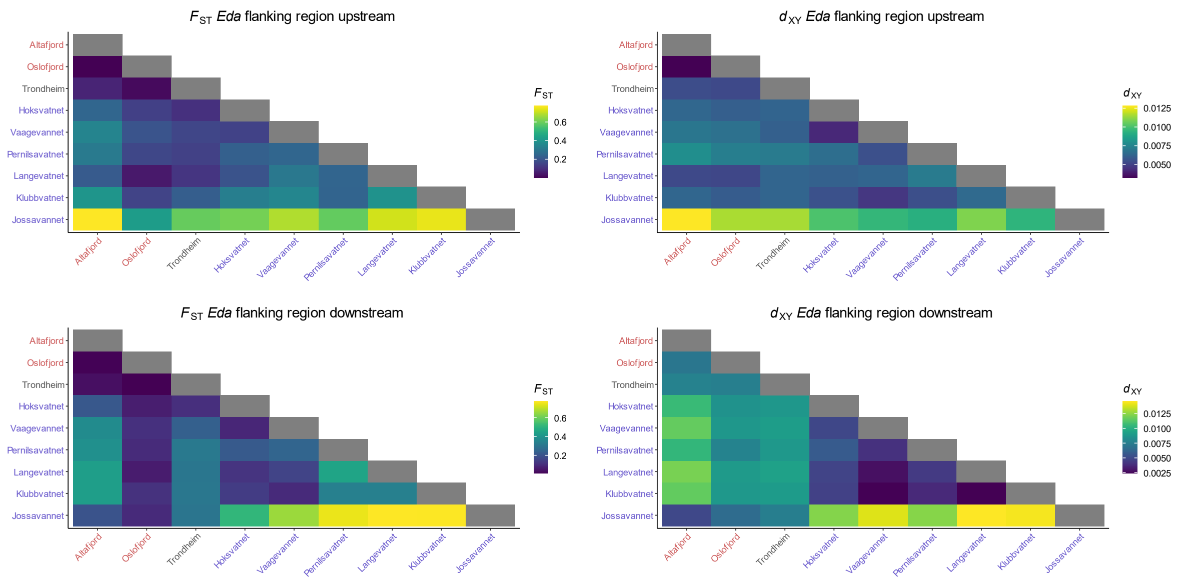


**Figure S3**: Window-based $F_{ST}$ and $d_{XY}$ statistics for all population pairs averaged for upstream (13 genomic windows) and downstream flanking regions (34 genomic windows) as a heatmap. Marine populations are written in red and freshwater populations in blue.

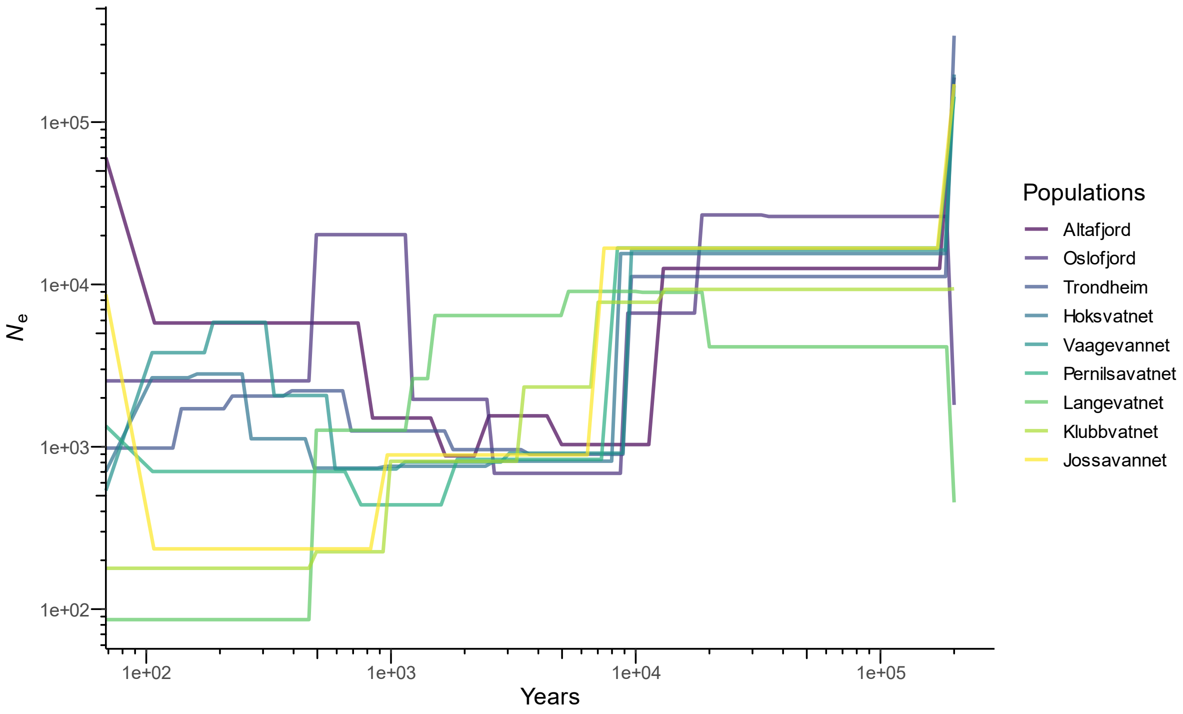


**Figure S4**: Demographic history for each population inferred with SMC++ using a generation time of 2 years and a mutation rate of 3.7 × $10^{-8}$ per site per generation.

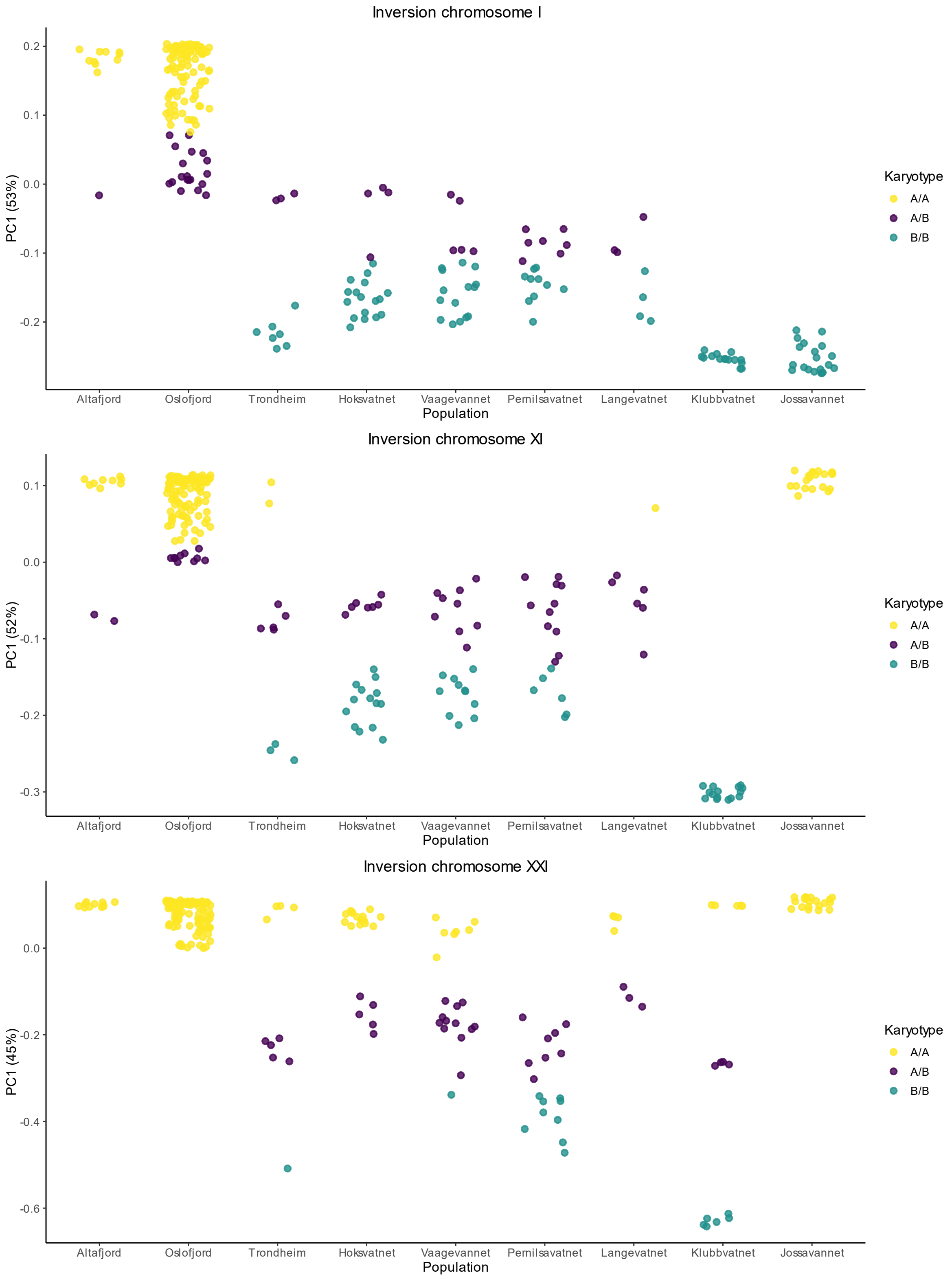


**Figure S5**: Clustering of the first principal component (PC1) for each sample in a population for each inversion. Genotype clusters are assigned based on *k*-means clustering with *K* = 3.

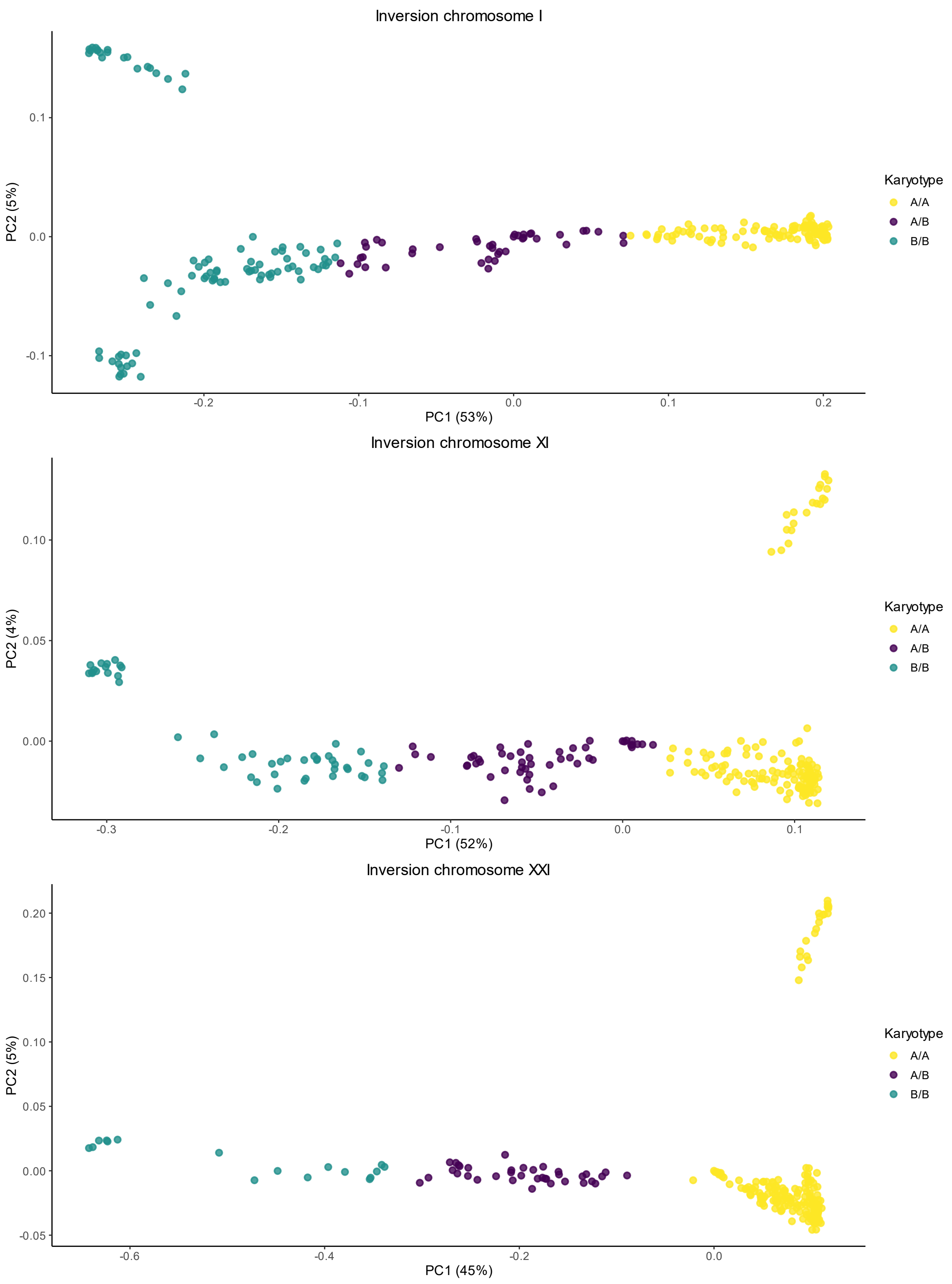


**Figure S6**: PCA for each inversion. Genotype clusters are assigned based on *k*-means clustering with *K* = 3.

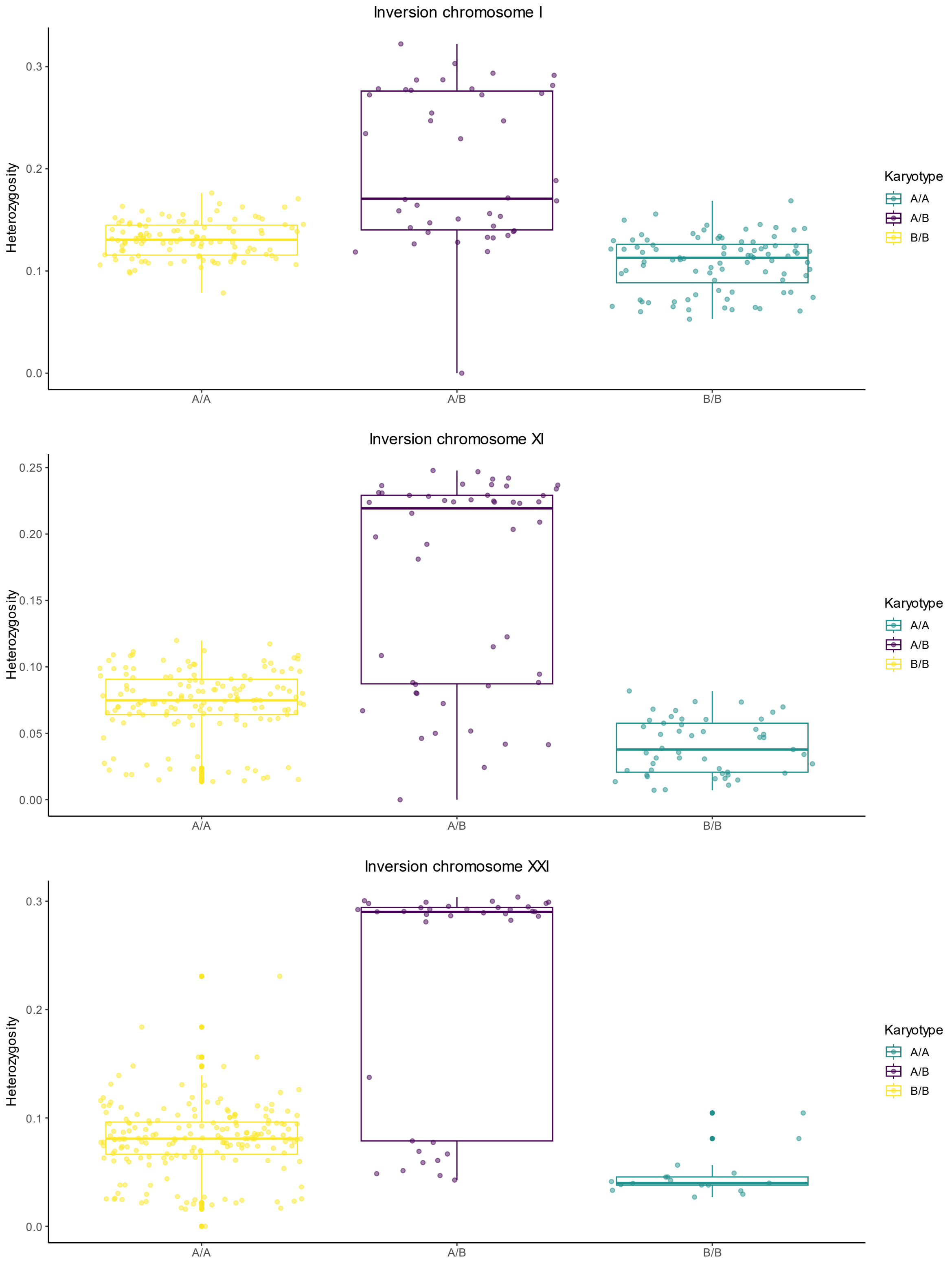


**Figure S7**: Proportion of heterozygosity for each classified genotype for each inversion. Genotype clusters are assigned based on *k*-means clustering with $K = 3$.

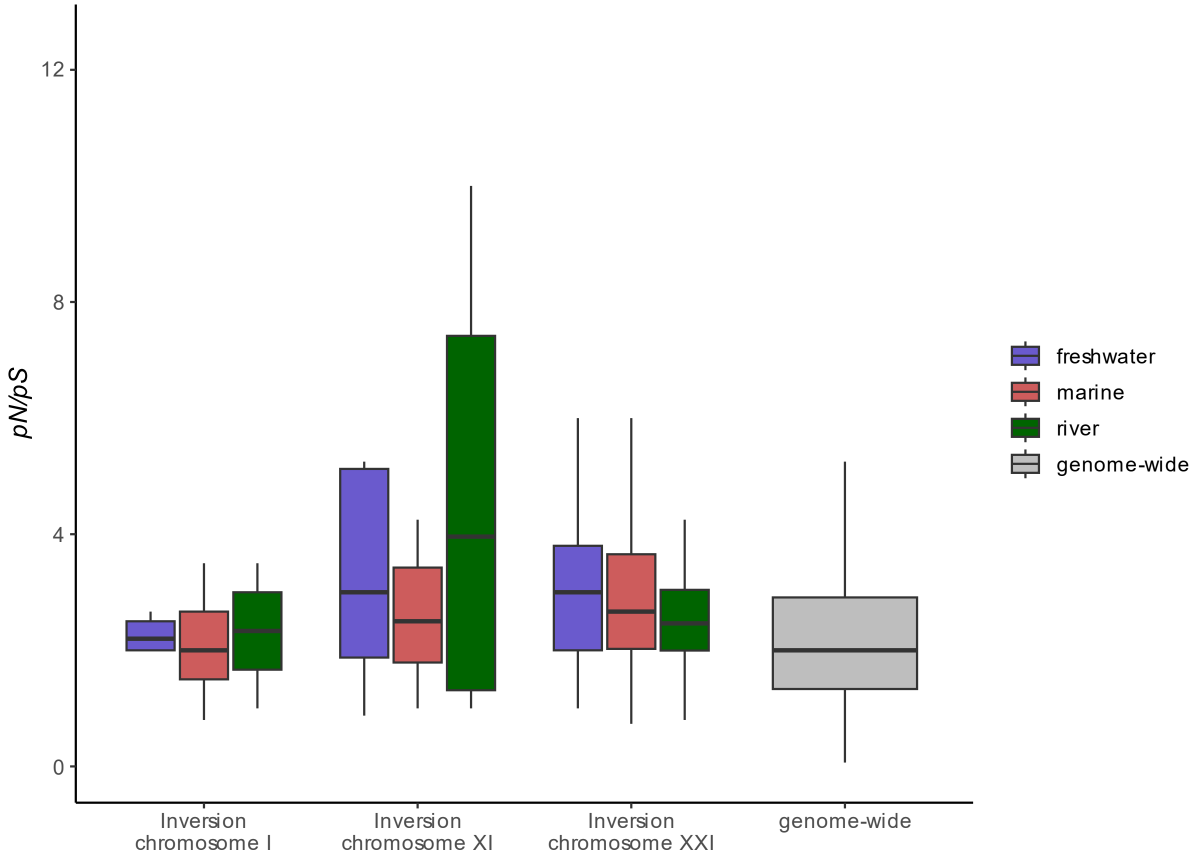


**Figure S8:** Ratio of nonsynonymous to synonymous sites (*pN*/*pS*) for the inversions on chromosome I, XI, and XXI for each group (marine, freshwater, and river) and genome-wide (excluding inversions) computed in 100 kb windows. The two-sided *t*-test showed no significant differences between the groups for each inversion or between a group within an inversion and the genome-wide *pN*/*pS*.

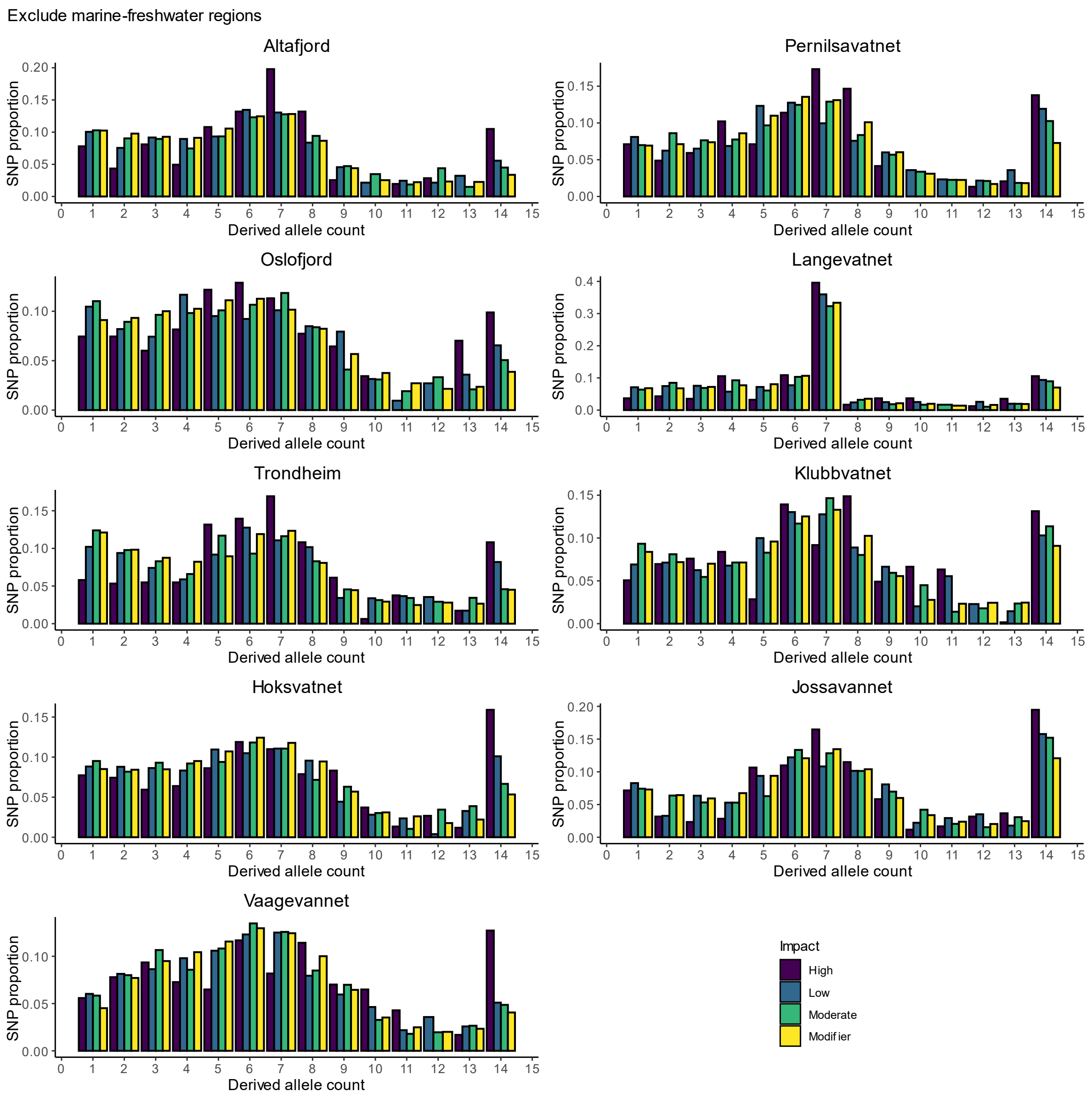


**Figure S9**: Derived site frequency spectra for the entire genome for each population for modifier, low, moderate, and high deleterious impact mutations (excluding the marine-freshwater divergent regions). Downsampled to the smallest diploid population size. Derived allele count of 0 is not shown.

#### Supplementary Tables

**Table S1.** Overview of the sample locations.

| Location | Latitude | Longitude | Habitat type | Lake age |
|---|---|---|---|---|
| Altafjord | 70.45298 | 23.77689 | Marine | / |
| Oslofjord | 59.66548 | 10.58540 | Marine | / |
| Trondheim | 63.42812 | 10.37346 | River | / |
| Hoksvatnet | 58.12242 | 8.10046 | Freshwater | 1,000 |
| Vaagevannet | 68.43912 | 16.15415 | Freshwater | 1,600 |
| Pernilsavatnet | 68.50971 | 15.88204 | Freshwater | 4,700 |
| Langevatnet | 68.37542 | 15.39080 | Freshwater | 9,600 |
| Klubbvatnet | 70.46198 | 23.79239 | Freshwater | 11,500 |
| Jossavannet | 70.60829 | 23.62477 | Freshwater | 12,900 |

**Table S2 (separate file)**. Sequencing results for whole-genome data for the total trimmed and mapped reads and the read coverage.
